# Distinct positional identity at the center of the caudal fin establishes forked shape

**DOI:** 10.64898/2026.05.16.725681

**Authors:** Eric Surette, John Gablemann, Katherine Backus, Brendan Fitzgerald, Tho Nguyen, Deirdre McKenna, Carmen Sofia Uribe Calampa, Sarah K. McMenamin

**Affiliations:** Boston College, Chestnut Hill MA 02467

## Abstract

The morphogenesis of complex vertebrate appendages requires precise regulation of growth, governed by distinct positional identities. The zebrafish caudal fin achieves a symmetrical, forked morphology through the regional specialization of the bony rays: peripheral rays are composed of relatively long, thick segments; while the central rays are made up of shorter, thinner segments, and their overall length is restricted. This length differential establishes the definitive forked shape of the organ. We asked whether these regional morphological differences reflect distinct underlying positional identities. Transcriptomic profiling of intact tissues from adult wild-type zebrafish suggested that central rays possess unique expression profiles, distinct from those of peripheral rays. We previously identified a treatment during embryogenesis that allows excess growth in the central rays, creating a truncate fin shape in adults–we asked whether this novel fin shape was caused by a peripheralization of the central rays. Indeed, the central rays of truncate fins were not only longer, but were composed of longer and thicker individual segments, reminiscent of peripheral rays. Further, gene expression in the central regions of truncate backgrounds showed signatures of peripheral identity. During development of the truncate phenotype, peripheral markers became expressed in more central domains of the growing truncate caudal fin, and in the supportive endoskeleton, the central hypural diastema was lost from the earliest stages. Ultimately, our results demonstrate how adult morphologies may be altered by shifts in positional identities. These findings clarify the anatomical patterning and molecular profiles that underlie regional specialization during caudal fin development.

## Introduction

Appendage function relies on precise morphological patterning, yet the developmental mechanisms that establish these patterns and positional identities remain incompletely understood. The caudal fin is a tractable vertebrate appendage with a straightforward anatomy. The external skeleton of the caudal fin consists of 18 bony rays composed of joints. The shape of the external caudal fin is defined by the relative lengths of these rays: in zebrafish, the peripheral rays are longer than the central rays, establishing a robustly forked shape (1,2). The internal base of the fin is supported by a series of modified caudal vertebrae that form the hypural complex. The central hypurals are separated by a diastema aligned with the body axis, which separates the upper and lower lobes of the fin (3–5).

Across teleost species, the caudal fin exhibits a range of morphologies, and caudal fin shape corresponds with hydrodynamic performance and specialization (6,7). While zebrafish possess a definitive forked caudal fin (with the shortest rays at the center) other species exhibit a spectrum of caudal morphologies, ranging from lunate and deeply forked, to truncate or rounded (with the longest rays at the center) (3–5,8).

We previously demonstrated that a transient pulse of *sonic hedgehog a* (*shha*) overexpression during late stages of embryonic development can repattern the shape of the adult caudal fin (9). Embryos exposed to *shha* pulse develop with central rays that show a loss of growth restriction, exhibiting higher proliferation and faster growth rates to reach lengths similar to those of the peripheral rays, and producing a triangular, truncate fin shape (see **Fig 1A-B**). The peripheral rays in these truncate fins are similar in length to the peripheral rays in forked fins of wild-type (WT) backgrounds. Truncate fins regenerate with relatively lengthened central rays to reestablish a forked morphology, suggesting a permanent shift in tissue identity relative to WT (9). Here, we asked if the shift in external shape and positional memory reflects the loss of a central identity, and adoption by the rays at the fin center of peripheral morphologies and molecular characteristics.

**Fig. 1:**
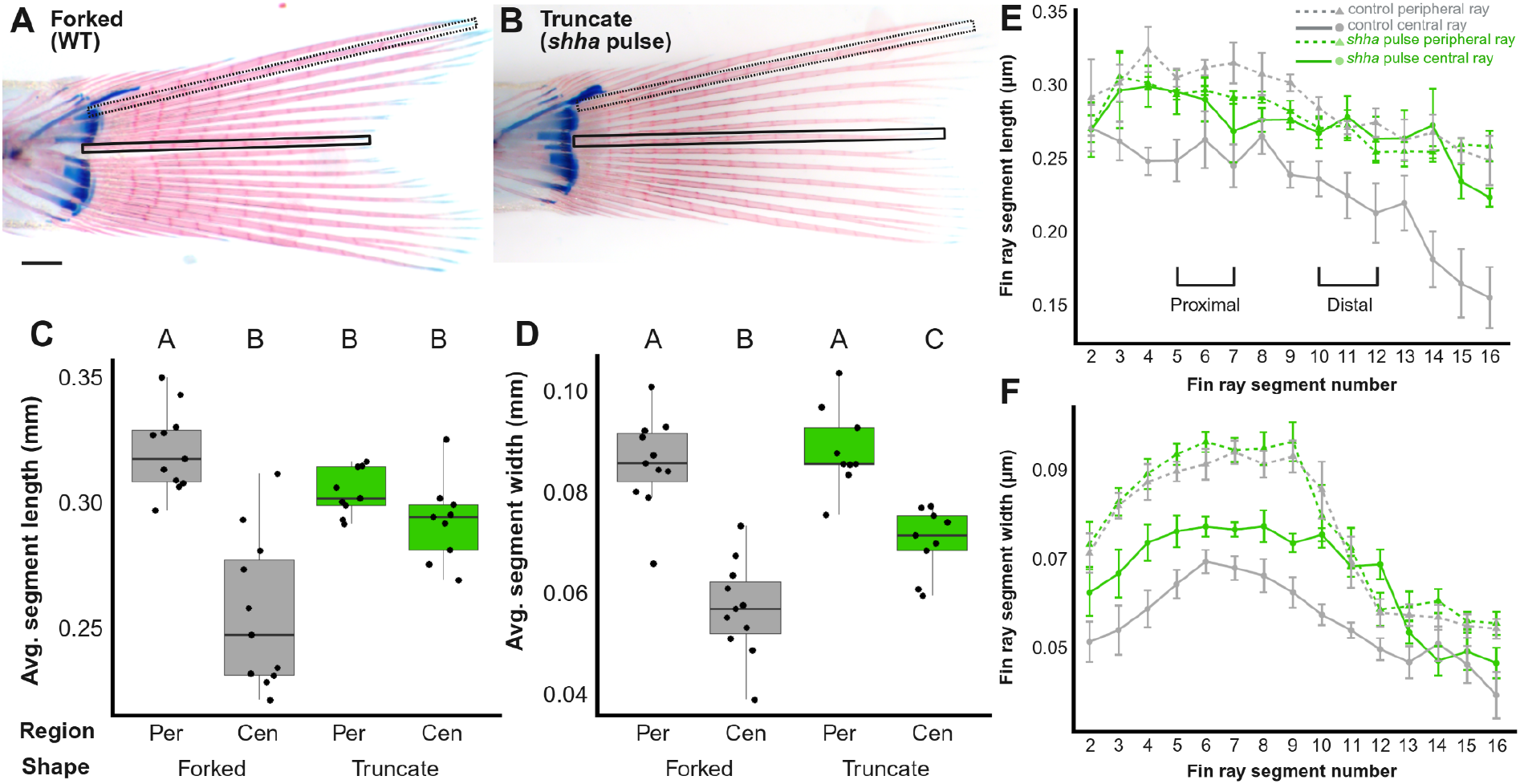
Central fin rays show distinct morphologies that are lost in truncate fins. (A–B) Representative images of zebrafish caudal fins exhibiting forked (A) and truncate (B) morphologies. Outlines indicate the central (solid line) and peripheral (dotted line) principal fin rays analyzed. Scale bar: 500 µm. (*C-D*) Quantification of fin ray segment length (C) and width (D) across different fin shapes and regional domains. Statistical significance was determined by a two-way ANOVA followed by Tukey’s post hoc test; groups sharing the same letter are statistically indistinguishable (p > 0.05). (*E-F*) Spatial profiles of segment lengths (E) and widths (F) for peripheral (dotted lines) and central (solid lines) rays, plotted by individual segment position along the proximodistal axis in forked (gray) and truncate (green) fins.

## Results

### Elongated central rays adopt peripheral segment morphology

In WT forked fins, the shortest central rays are composed of bony segments that are thinner and shorter than those in the periphery **(Fig. 1A)**. We asked whether the overgrowth of central rays induced by a transient embryonic pulse of *shha* (9) was accompanied by a shift in this underlying segment morphology. Indeed, the central rays of truncate fins were not only longer, but their individual segments grow longer and thicker, closely resembling the morphology of peripheral ray segments **(Fig 1B-F)**. The peripheral rays from both backgrounds showed similar morphologies, suggesting that this shift did not reflect a universal change in ray proportions, but rather shifts specific to the central rays (**Fig 1C-F**). Location of ray bifurcations remained unchanged between forked and truncate backgrounds **(Fig. S1)**.

Adult fin shape is imprinted into the fin primordium during embryonic development, and transiently increasing Shh activity during this period increases the growth potential of the central rays that will develop (9). We asked if other disruptions during this critical developmental window could induce analogous phenotypic shifts. Like Shh, the Wnt pathway is crucial in appendage patterning (10), but we found that increasing or antagonising the Wnt pathway during the late stages of embryogenesis had no effect on adult fin morphology **(Fig. S2)**. In contrast, larvae in which the ventral fin fold was amputated prior to ray ossification (2,13) developed into adults with externally truncate fins, capable of regenerating with a truncate shape **(Fig. S3, S4)**. However, since treatment with *shha* pulse produced a more consistent phenotype, we used *shha* pulse to induce truncate phenotypes for the following analyses.

### Peripheral tissues encroach into the central regions of developing truncate fins

Given that mature central rays adopt peripheralized morphological characteristics following an *shha* pulse, we asked whether this represents an expansion of peripheral developmental programs at the expense of central boundaries during early fin patterning. The transcription factor *alx4a* is associated with anterior identity in developing paired appendages (14,15); accordingly, developing forked fins show restricted *alx4a* domains in the areas that will form the peripheral-most regions of the fin (**Fig 2A;** (2)). In developing truncate fins, the posterior domain of *alx4a* expands into the fin center, decreasing the separation of the two domains (**Fig. 2B-C**).

**Fig. 2:**
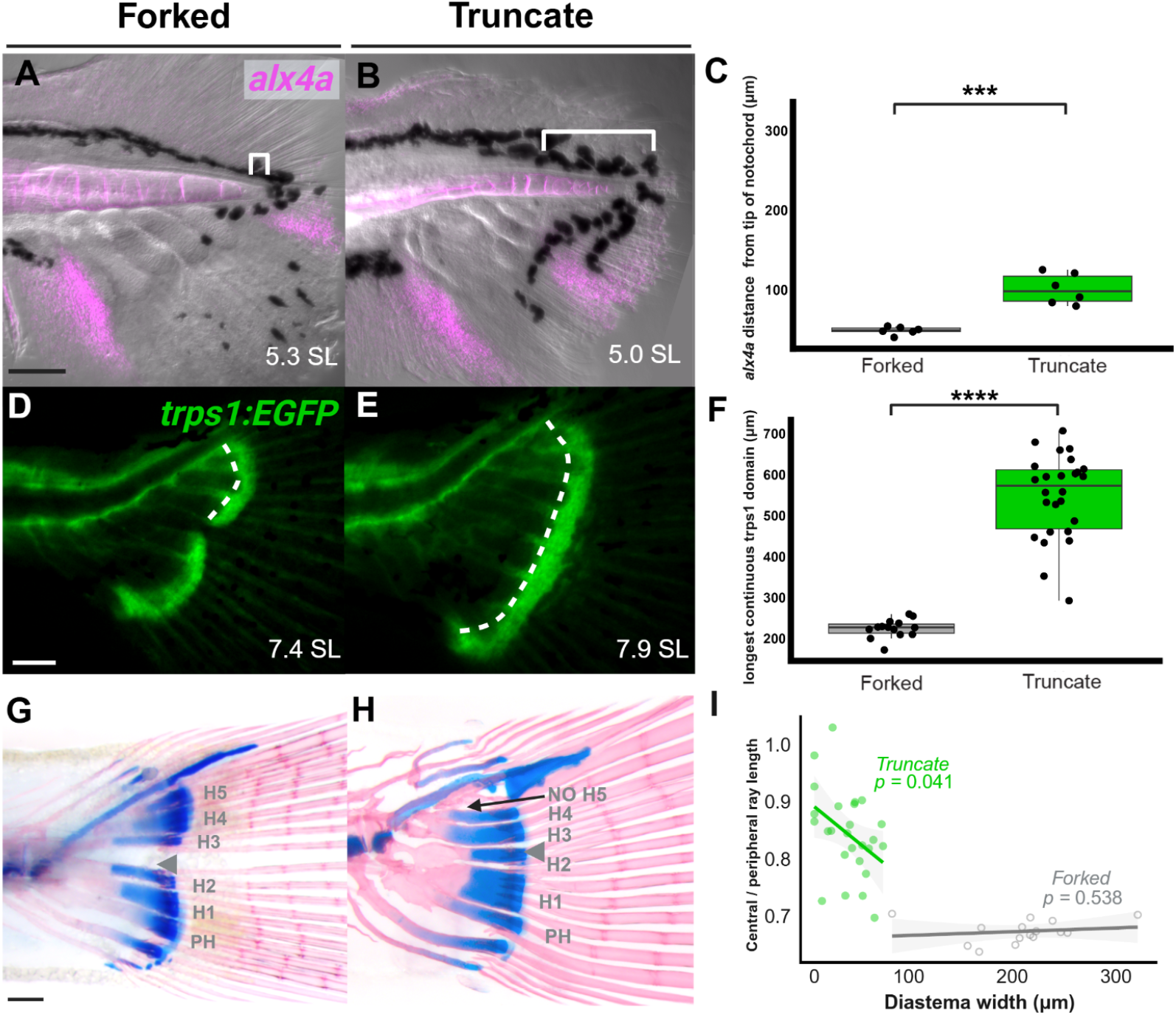
Central features are absent in truncate fin development. (*A-B*) Domains of *alx4a* and in the larval forked fin (*A*) and truncate (*B*) caudal fin at 10 dpf. (*C*) Size of posterior *alx4a* domain, as measured from the tip of the notochord (shown in brackets in A and B). Significance determined by Welch two-sample T-test. (*D, E*) *trps1:EGFP* labels the plates of connective tissue which connect the hypural complex to the fin rays. Dotted white lines indicate measurement distance; both individuals imaged at 13 dpf. (*F*) Quantification of the longest continuous trps1 domain, starting at the location closest to the posterior notochord. Significance determined by Welch two-sample T-test. (*G*) The zebrafish ural endoskeleton consists of five hypurals (H1-5), separated by the hypural diastema (gray arrowhead). (*H*) Truncate fins fail to form a hypural diastema and exhibit hypural defects. Scale bar: 200 µm. (*I*) Caudal fin shape truncate fins correlates with the size of the residual hypural diastema. Significance of relationship between ‘central / peripheral fin ray length’ and hypural diastema size determined by linear-mixed effects model. Scale bars: 100 µm: (*A-B*), (*E-F*); 200 µm: (*G-H*).

The supportive hypurals develop in close association with connective tissues in which *trps1:eGFP* is active (2); these connective tissue plates show clear separation at the hypural diastema **(Fig. 2D)**. In contrast, truncate fins developed with a single continuous plate, and the central diastema region is absent **(Fig. 2E-F)**. (We note that larval fin fold amputation results in a smaller space between the *trps1:eGFP* domains, although the plates still appear separate, **Fig. S4G-H)**.

### Caudal fin endoskeletal morphology and patterning correlates with external fin shape

The endoskeleton of WT forked fins shows a clear diastema between the central hypurals **(Fig 2G)**. The failure of the *trps1* domains to form a distinct separation **(Fig. 2E)** presages a collapse of the hypural diastema in the adult truncate fins **(Fig. 2H)**. The hypural complex in truncate backgrounds showed a range of aberrant phenotypes, including fusion between hypurals, loss of hypurals and formation of supernumerary hypurals (**Fig. S5B-G**). Truncate fin complexes showed a number of expanded peripheral features, specifically an enlarged opistural cartilage (**Fig. S5H**), increased numbers of epurals **(Fig. S5J)** and an increased number of procurrent rays **(Fig. S5I)**. While vertebrae and preural spine numbers remained unchanged in a truncate background **(Fig. S6**), the notochord showed reduced flexion and the hypural complex misaligned with the body axis (**Fig. S7**). These data suggest the internal caudal fin skeleton expands concomitantly with the loss of the diastema, without altering preural caudal fin patterning.

We asked if changes in shape and fin ray identity correlated with the degree of diastema loss, we analyzed the morphology of the caudal fin in relation to the diastema. In truncate fins, fish with smaller diastemas predict longer central rays in truncate backgrounds (**Fig. 2I**). Diastema size was also inversely correlated with central ray segment morphology in truncate fins, while peripheral segment morphology showed no correlation (**Fig. S8**). Together, these results indicate that the loss of the hypural diastema corresponds with a shift toward peripheral morphologies at the center of the fin.

### The central regions of truncate fins adopt peripheral patterns of expression

To determine if the morphological differences between the center and the periphery of the WT fin reflect underlying molecular differences, we analyzed their regional transcriptomes. We identified 81 genes that are differentially expressed between wild-type central and peripheral rays, which we refer to as region-specific transcripts **(Fig. 3A-B, Fig. S9A-B)**. Gene Set Enrichment Analysis suggests that these differences may be driven by discrete gene regulatory networks, involved in disparate biological processes **(Fig. 3C)**.

**Fig. 3:**
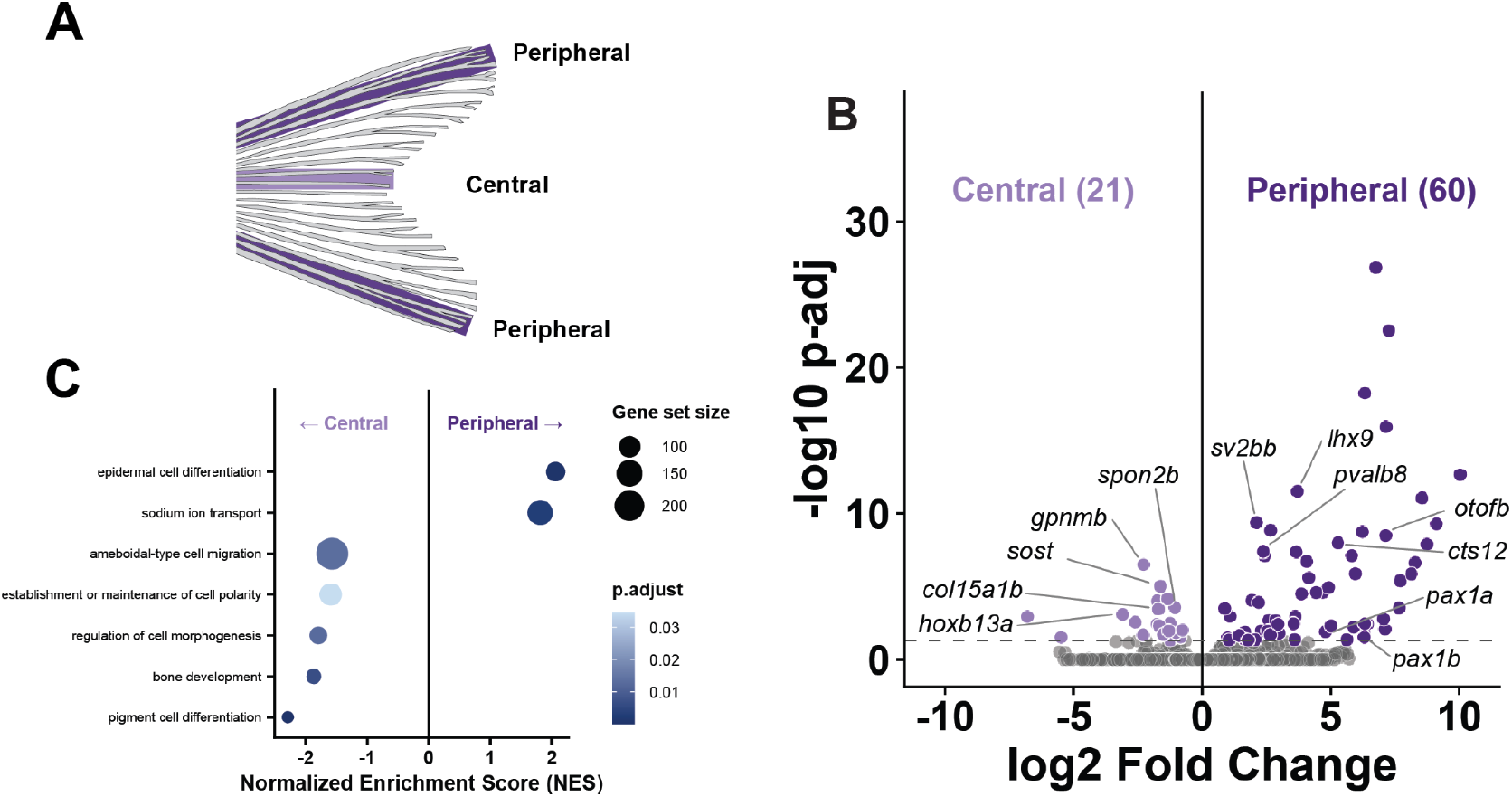
Central and peripheral rays of forked fins show distinct transcriptomic profiles. *(A)* Central (light purple) and peripheral (dark purple) regions extracted from WT forked fins for bulk RNA sequencing. *(B)* Comparison of transcripts from central and peripheral rays reveals differentially expressed genes. (*C*) GSEA analysis showing developmental terms that are significantly enriched in either central or peripheral rays.

We asked if morphologically “peripheralized” truncate central rays molecular signatures characteristic of central rays in WT forked backgrounds. Indeed, in truncate backgrounds, peripheral and central regions were more similar in expression than peripheral and central regions from WT forked backgrounds **(Fig 4B)**. Moreover, the central regions from a truncate background showed more similarity to the peripheral regions than the central regions from a WT forked background **(Fig 4B)**. Rather than maintaining their unique wild-type signature, truncate central rays undergo a broad molecular convergence, adopting a transcriptomic profile that quantitatively mirrors and clusters with the peripheral samples (**Fig. 4B-C**).

**Fig. 4:**
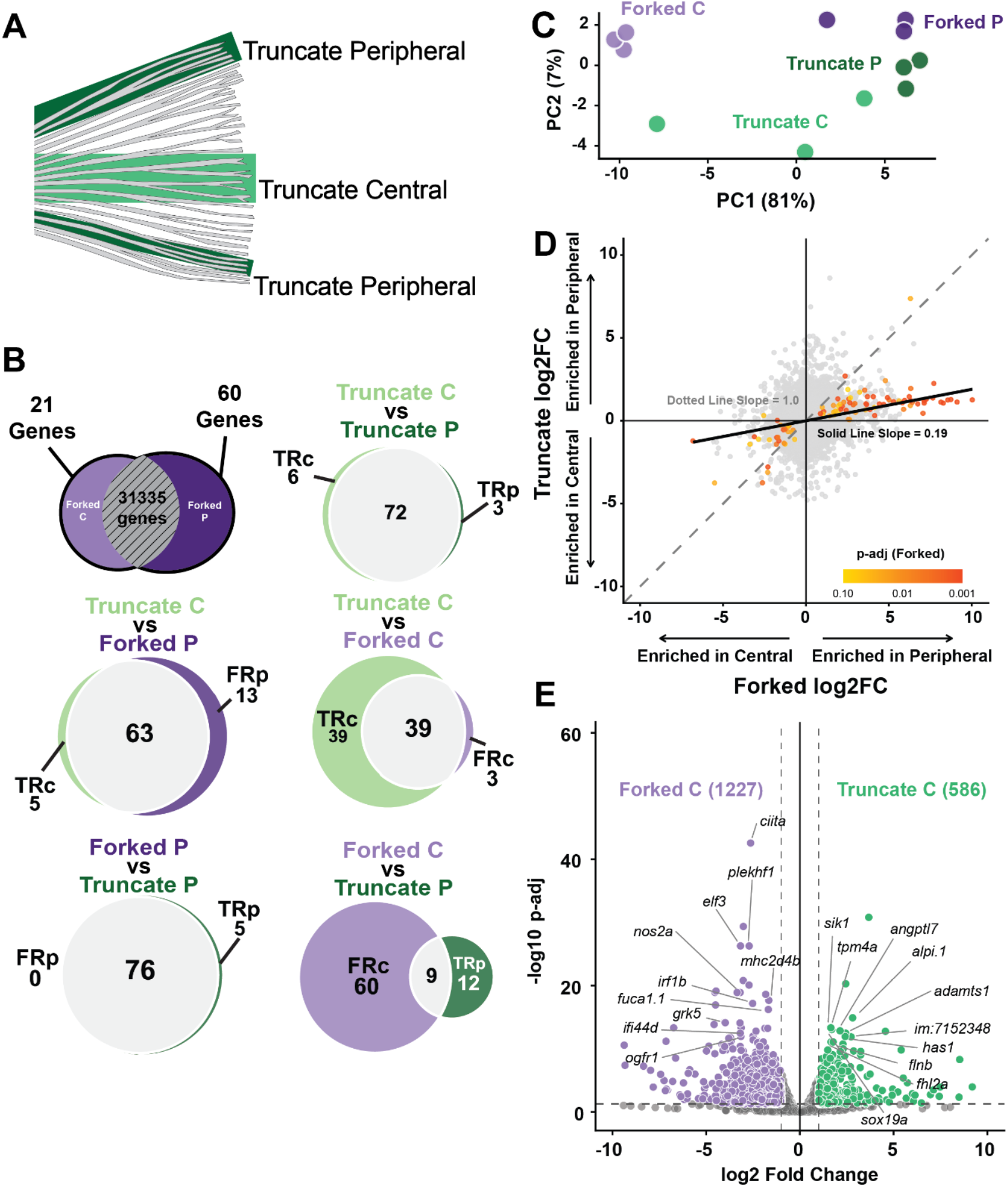
Central rays of truncate fins adopt a peripheral molecular profile. (A) Central (light green) and peripheral (dark green) regions extracted from truncate fins for bulk RNA sequencing. (*B*) Venn diagrams showing comparisons of enrichment for the 81 region-specific genes. The number in gray represents genes that are expressed at a similar level in both conditions; a higher gray number indicates that the two conditions are more similar *(C)* Log fold change plot showing central-peripheral transcriptomic identity of each gene in both WT forked and truncate fins. Points that achieved a *padj* value below 0.10 are colored, with warmer colors representing a lower *padj* value. The dotted line (slope = 1) represents equivalent expression in both WT forked and truncate fins, and the black trendline represents the line of best fit for all points, demonstrating a shift towards central and peripheral similarity in truncate fins. *(D)* A PCA plot made using the 81 region-specific genes. (*E*) Volcano plot representing genes that are differentially expressed in WT forked and truncate central rays.

To evaluate this convergence at the individual gene level, we examined the central-to-peripheral log2 fold changes across both genetic backgrounds **(Fig. 4D)**. The resulting line of best fit significantly deviated from the 1:1 equivalence line which denotes equal expression in both conditions. This deviation demonstrates a shift toward transcriptomic similarity between central and peripheral domains in truncate fins **(Fig. 4D)**. Consistent with this convergence, direct differential expression analysis between the central and peripheral rays within truncate fins confirms a striking loss of regional uniqueness (**Fig. S10A**).

We examined how this collapse of regionality was manifested across specific gene cohorts. While comparing truncate and forked samples highlights broad global differences in gene expression and functional enrichment between wild-type and truncate fins overall (**Fig. S11A-B**), a direct comparison specifically between the central domains of forked and truncate fins reveals a targeted transition in the regional molecular profile **(Fig. 4E)**. Pairwise differential expression analysis between these central regions identified 1,227 transcripts with significantly enriched expression in forked central rays relative to truncate central rays, representing a widespread loss of the baseline central-specific signature in the truncate background **(Fig. 4E)**. This comparison highlights candidate genes that are differentially expressed and likely drive this localized shift toward a peripheral identity (**Fig. 4E**). These findings show that the truncate phenotype is characterized by a disruption of normal central molecular and morphological identity, accompanied by localized overgrowth of the central rays.

## Discussion

Positional identity can be inferred by expression and morphology (15–20). The definitive forked shape of the zebrafish caudal fin reflects regional differences in position-specific growth of the fin rays. Central rays likely possess specific positional information that inhibits growth and widening of growing segments, and limiting their ultimate lengths relative to peripheral rays. Disrupting the positional identities that imprint this local growth rate can cause the central rays to adopt a more peripheral identity. Brief overactivity of *shha* in the fin tissues or early amputation of the larval fin fold disrupt processes by which these fates are imprinted, altering the identities that regulate growth throughout larval, juvenile and adult development. Each of the 18 rays may have a unique positional identity. We have focused on the characteristics of the peripheral-most and central-most regions of the forked fin, and it is likely that positional identity exists in a spectrum across these individual rays from central to peripheral.

*alx4a* is expressed in anterior regions of the dorsal and anal fins, and is expressed bilaterally in both the upper and lower peripheral regions of the developing caudal fin (2). In a developing truncate background, these peripheral domains collapsed closer to one another, and the *alx4* domain in the posterior region (fated to become the upper lobe) grew to cover a large portion of the fin primordium. Nonetheless, there was still a space between the *alx4* domains. In contrast, in a truncate background, there was no space maintained between the *trps1* domains of connective tissue. Either as a cause or an effect of this single connective tissue domain, the hypurals in a truncate background developed with no separating diastema–the loss of a central character.

In adult WT, the central and peripheral regions of the uninjured forked fin show distinct patterns of expression. Peripheral regions show enriched expression of critical skeletal patterning factors, including *lhx9* and *pax1a/b*, which are known regulators of fin bud development and chondrocyte differentiation **(Fig 3B, Fig. S9A-B;**(21–24)). We also identified peripheral enrichment of ion channels and transporters, such as *chrna7a* and *slc6a13*. This provides a direct molecular basis for the localized bioelectric pathways that are known to drive zebrafish fin allometry and scaling (25,26). As we did not separate the upper and lower peripheral regions of the fins in this analysis, we factors enriched in the pooled peripheral tissue may be elevated in one or both peripheral regions.

The central region is characterized by elevated expression of posterior Hox factors including *hoxb13a*, which is required for early caudal fin patterning, expressed in the posterior and central regions of the developing fin (1,9). Also enriched in the central regions was *sost*, which encodes an inhibitor of the Wnt and BMP pathways and is likely to inhibit bone growth (27). Enriched transcripts in the central fins likely reflect and mediate a central identity, regulating and limiting the growth of the central rays to create the forked shape of a WT zebrafish fin.

Certain disruptions to early fin patterning appear to abolish the central identity, with rays at the center of the fin adopting morphological characteristics and expression patterns more appropriate for peripheral rays. This could be interpreted as the prevalence of peripheral over central identities: in the absence of central signals, peripheral identities take prevalence.

Posterior Hox genes tend to take prevalence over more anterior Hox factors in determining positional identities and phenotypes in body segments (28,29), and apparent posterior dominance has been observed during limb regeneration (30). In the absence of growth-inhibiting signals at the center of the fin, central rays default to a more peripheral identity and are permitted to grow longer.

The hypural diastema serves as a key morphological marker of central identity in the teleost caudal fin, developing simultaneously with the first principal rays and the structural separation of connective tissue (3,7). In basal actinopterygians lacking a hypural diastema, the connective tissue of the hypurals typically forms a uniform plate (3,7). Truncate fins show a regional uniformity across the caudal fin connective tissue plates and rays that is reminiscent of certain more basal actinopterygians. We propose that the emergence of a distinct central identity provided a developmental mechanism to actively limit the growth of central rays, effectively decoupling the fin center from the periphery. This localized growth restriction may have permitted further fin specializations, as central and peripheral regions could be independently modulated to adopt different relative growth rates.

Our work suggests that underlying the forked caudal fin shape are distinct morphological and transcriptomic identities in the central and peripheral rays. These positional identities are imprinted during embryogenesis, and disruptions to early patterning can shift in the regional identities: the hypural diastema collapses and rays at the center of the organ adopt peripheral characteristics and identities. The positional identities that inform shape can be modulated to generate morphological diversity.

## Materials and Methods

### Animal husbandry

All experiments with zebrafish were done in accordance with protocols approved by the Boston College Institutional Animal Care and Use Committee. Zebrafish were reared under standard conditions at 28°C with a 14:10 light:dark cycle. Fish were fed marine rotifers, *Artemia*, Adult Zebrafish Diet (Zeigler, Gardners PA, USA) and Gemma Micro (Skretting, Stavanger, NOR).

For developmental serial imaging, siblings were reared in individual containers so individuals could be identified and tracked. The *shha* pulse causes changes in early growth rate, so size-match control individuals were used for comparisons; standard length (SL) is reported throughout. Note that prior to development of the hypural complex, notochord length was measured, and is referred to as SL, per (13).

### Transgenic and mutant lines

Transgenic lines used were *Tg*(*hsp70l:shha-EGFP*)(31) to induce *shha* pulse (below), *Tg(sp7:GFP)b1212* (32) to visualize osteoblasts, *Tg(p7*.*2sox10:mRFP)* (33) to visualize chondrocytes, *Tg(trps1:EGFP)* ^*j127aGt*^ (34) to visualize the endoskeletal connective plates, and *flk1:DsRed2* (also called *Tg(kdrl:dsRed2)*(35)) to visualize blood vessels.

### Imaging

Zebrafish were anesthetized with tricaine (MS-222, ∼0.02% w/v in fish system water). Live anesthetized or cleared and stained (36) or anesthetized individuals were imaged on an a Leica Thunder Imager Model Organism using a sCMOS monochrome camera, a Zeiss AxioImager Z2 using a Hamamatsu Flash4.0 V3 sCMOS camera, or a Leica IVESTA 3. Identical exposure times and settings were used to compare experimental treatments and capture repeated images of fins. Images were correspondingly adjusted for contrast, brightness and color balance using FIJI (37), and compiled using BioRender.

### *shha* pulse to generate truncate fins

We induced *shha* overexpression pulse in embryos using our previously developed protocol (9). Sibling larvae treated with heat shock but were negative for GFP were kept as negative controls, and typically embryos with the brightest GFP were kept as the *shha* pulse group.

### Drug treatments and larval fin fold amputations

Sibling groups of fish were treated for 24 hours with different concentrations of the Wnt agonist BIO or the Wnt antagonist IWR-1 (11,12). Following treatment, larvae were rinsed and allowed to grow under standard conditions. Larvae were anesthetized at 6 days post fertilization using a tricaine solution and placed on a dish prepared with 2% methylcellulose and larval water. The ventral larval fin fold was mechanically removed using a scalpel.

### Fin ray morphology quantifications

Caudal fins from size-matched *shha* pulse and control sibling fish were imaged. The 2nd dorsal ray was used as the representative peripheral ray, and the central-most rays were considered central–in fins with 18 rays the 9th ray was used as representative. Individual landmarks were placed using the R package StereoMorph (38) were placed along the proximal and distal ends of each bony segment to capture segment length, and at the dorsal and ventral sides of each segment at the segment midpoint to capture segment width. An additional landmark was placed at the ray bifurcation of each ray. As the first proximal segment was frequently obscured by scales, this segment was excluded from analyses.

### Bulk RNA sequencing

Intact caudal fin rays were extracted from adult sibling WT forked and truncate fins (>1 year of age). Fish were anesthetized and tweezers were used to extract the 2nd-most dorsal and ventral fin rays, and the two central-most fin rays. The 2nd-most dorsal and ventral rays were pooled to generate the “peripheral” sample. Rays were stored in Thermo Fisher RNAlater Stabilization Solution (Cat. #: AM7021), and shipped to Azenta Life Sciences for RNA extraction, library preparation and sequencing. Raw sequence reads were aligned to zebrafish GRCz11.114 and called using Ensembl GRCz11 gene annotation. Differential gene expression analyses were performed with DESeq2 (39). Genes were considered significantly expressed if they showed a log2 fold difference higher than 1 and a false discovery rate lower than 0.05. Gene Ontology analysis was performed using clusterProfiler (40), with a q-value cutoff of 0.01.

### Whole mount fluorescent *in situ* hybridization

Fish were anesthetized, fixed for 30 minutes in 4% PFA, and dehydrated then rehydrated in a methanol series. Fish were stained as described (41), with the modification that all 0.2x SSCT washes were only performed twice. RNAscope Multiplex Fluorescent Reagent Kit v2 (ACD Bio-techne, 323100) was used to target *alx4a;* ACD Bio-techne 1306151-C4.

### Statistical analysis

Analyses were performed in RStudio. Data were analyzed with Welch two-sample, two-tailed t-test, Wilcoxon signed-rank test, ANOVA followed by Tukey’s honest significant differences (using 95% family-wise confidence level), Fligner-Killeen test, or linear mixed-effects model followed by Tukey’s honest significant differences (using 95% family-wise confidence level). In graphs showing pairwise comparisons, significance is indicated as follows: * *p* < 0.05, ** *p* < 0.01, *** *p* < 0.001. Each plotted data point represents a single measurement from a single fish, unless otherwise noted in the figure legend.

## Acknowledgements

For fish care and assistance, we thank the BC Animal Care Facility and members of the McMenamin Lab past and present. For the development of our landmarking protocol, we thank Dr. Yinan Hu. For sharing fish lines, we thank Drs. González-Rosa, Harris, Karlstrom, North, Smeeton and their labs. For sample processing bulk RNA sequencing, we thank Azenta Life Sciences. For assistance and use of the Zeiss AxioImager Z2, we thank Bret Judson of the Boston College Imaging Facility. For statistical assistance and troubleshooting, we thank Dr. Melissa McTernan.

Funding provided through NIH MIRA R35GM146467

---

**Fig. S1:**
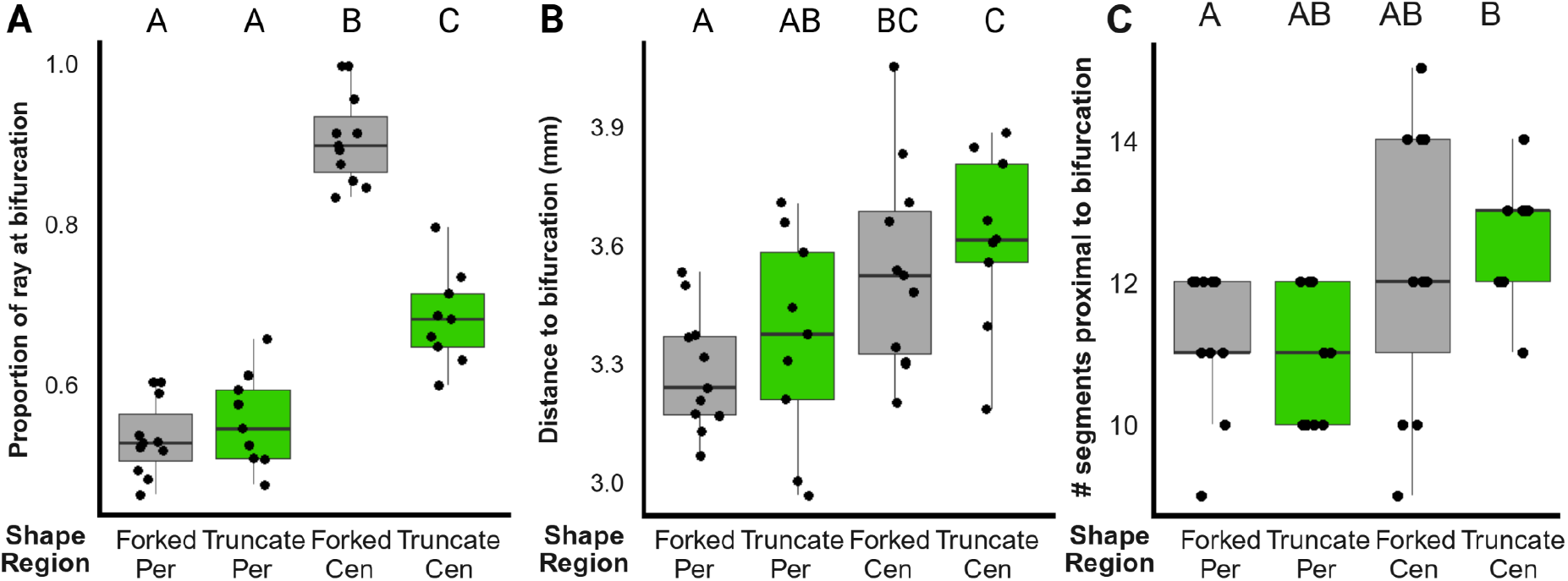
Individual rays form bifurcations at locations proportional to ray length. (*A*) The proportion of the overall ray length at which bifurcations form, (*B*) the absolute distance to bifurcations, and (C) the number of segments proximal to bifurcations. Significance determined by ANOVA followed by Tukey post hoc test. Statistically indistinguishable groups are shown with the same letter (threshold for significance *p* < 0.05).

**Fig. S2:**
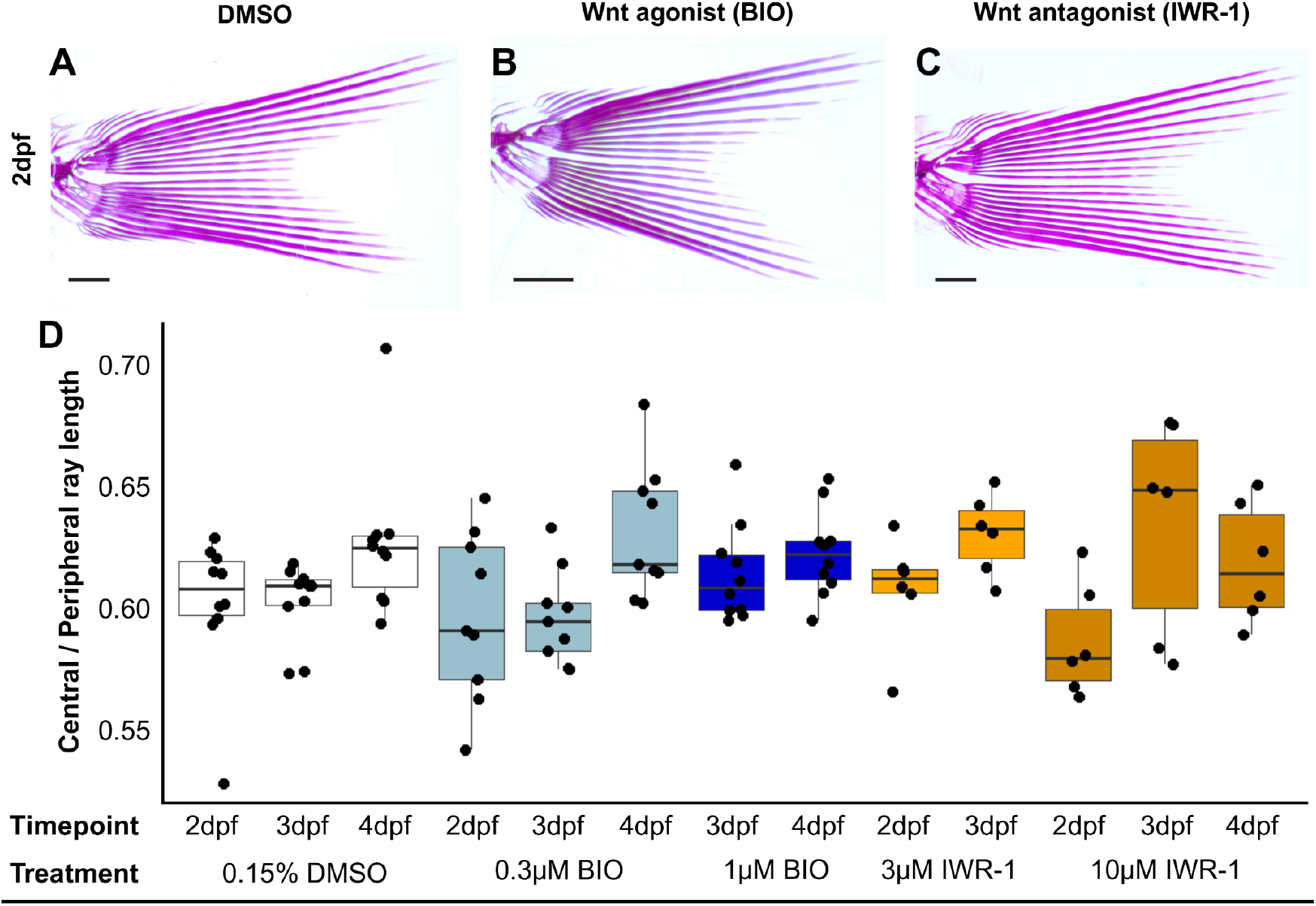
Modulating Wnt pathway during early development does not change adult fin shape. (*A-C*) Adult caudal fins following early treatment with Wnt agonist or antagonist. Scale bars: 200µm. (*D*) Caudal fin shape does not change following treatment with pharmacological agents affecting Wnt pathway. No groups are significantly different from another, as determined by two-way ANOVA followed by Tukey post hoc test.

**Fig. S3:**
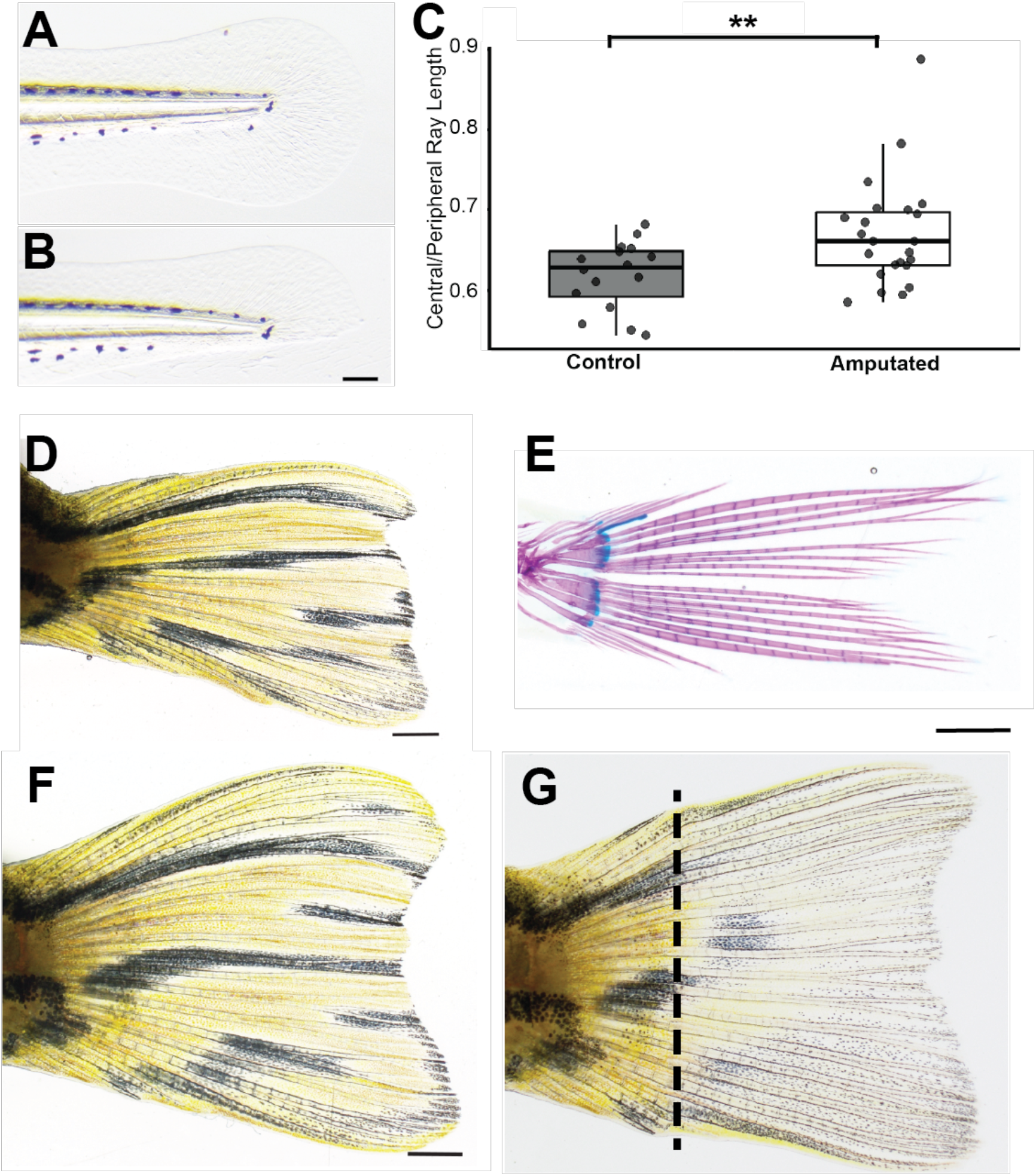
Larval fin fold amputation disrupts adult shape. *(A-B)* The zebrafish fin fold before (A) and after *(B)* ventral fin fold removal. Scale bar: 100 µM *(C)* Quantification of fin shape between sibling controls and individuals amputated at 6 days post fertilization. Significance determined using student’s t-test. *(D-E)* Adult caudal fins grown from amputated larval fin folds. Scale bar: 1 mm *(E). (F-G)* Adult fin following larval fin fold removal, before *(F)* and after *(G)* amputation and regeneration of adult tissues. Scale bar: 1 mm

**Fig. S4:**
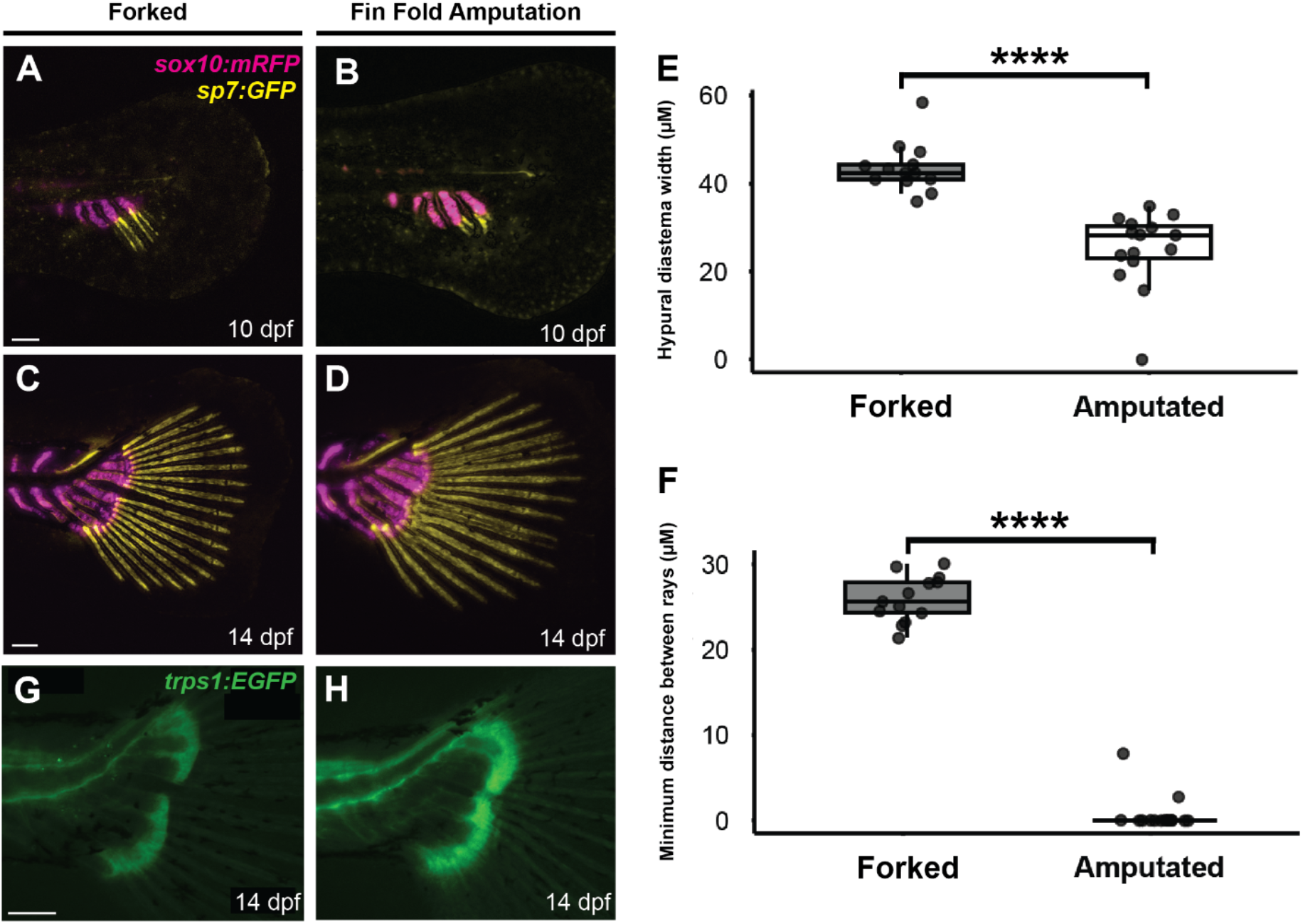
Larval fin fold removal disrupts early skeletogenesis. (*A-D*) Chondrocytes (*sox10;* magenta) and osteoblasts (*sp7;* yellow) in the developing uninjured WT caudal fin (*A, C*) and following fin fold amputation (*B, D*). Scale bar: 100 µM. (*E)* Quantification of hypural diastema width in 14 dpf zebrafish. Significance calculated using student’s t-test. *(F)* Quantification of the minimum distance between rays in 14 dpf zebrafish. Significance calculated using wilcox test. (*G-H*) *trps1:EGFP* expression in the connective tissue plates at the hypural-ray junction following larval fin fold amputation. Scale bar 100 µM.

**Fig. S5:**
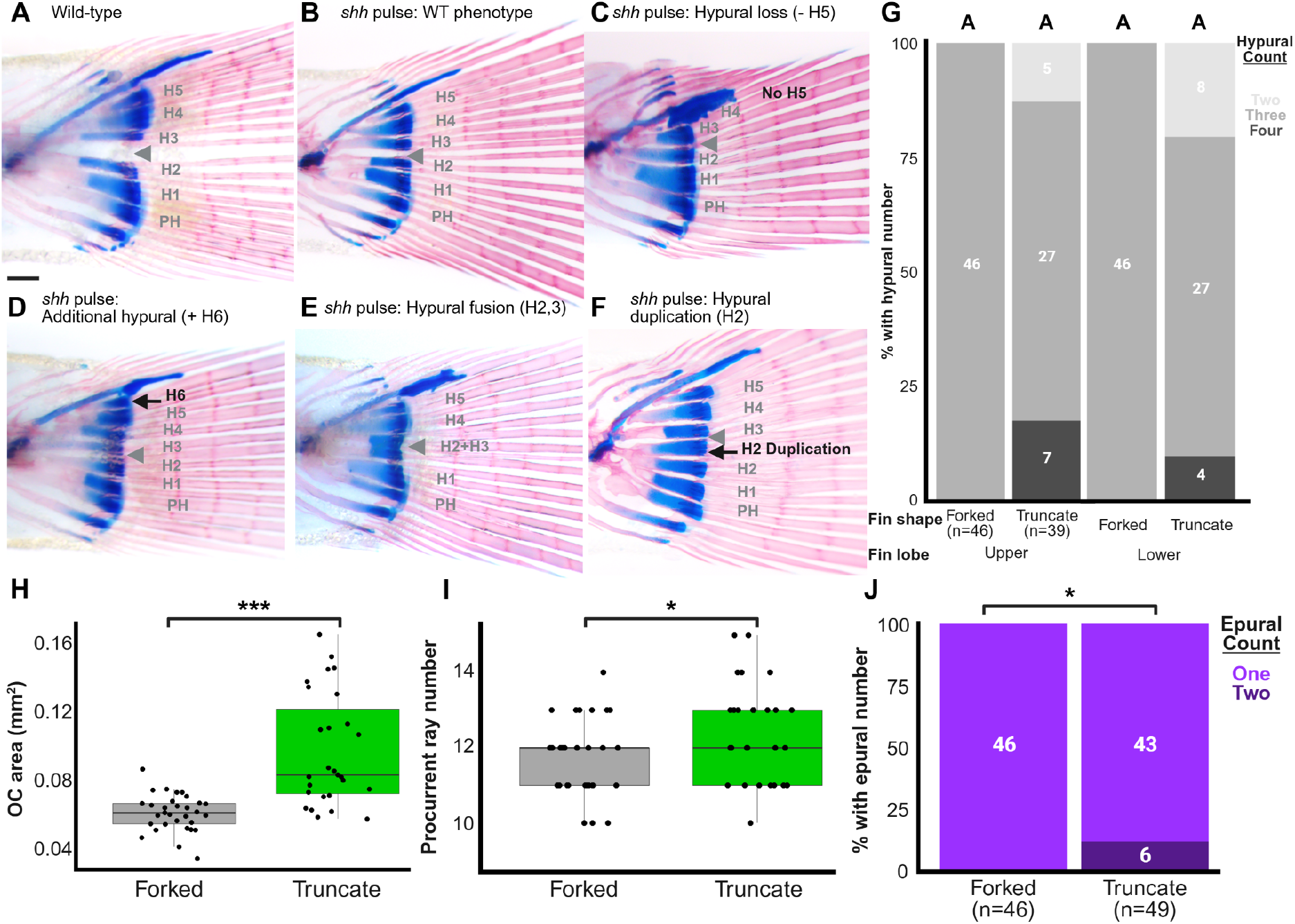
Hypural complex of truncate fins show variable anatomical malformations. (A) Hypural complex of a WT forked fin. (B-F) Variable hypural phenotypes of truncate fins, including hypural loss (*C*), gain (*D*), fusion (*E*) and branching (*F*). Scale bar: 200 µM. (*G*) There is no dorsoventral bias in the hypural aberrations; analyzed by two-way ANOVA followed by Tukey’s post hoc test. Statistically indistinguishable groups are shown with the same letter (threshold for significance *p* < 0.05). (*H-J*) Size of the opistural cartilage, total number of procurrent rays (*I*), and number of epurals (*J*). Significance by Welch two-sample T-test (*H-I*) or Wilcoxon signed-rank test (*J*).

**Fig. S6:**
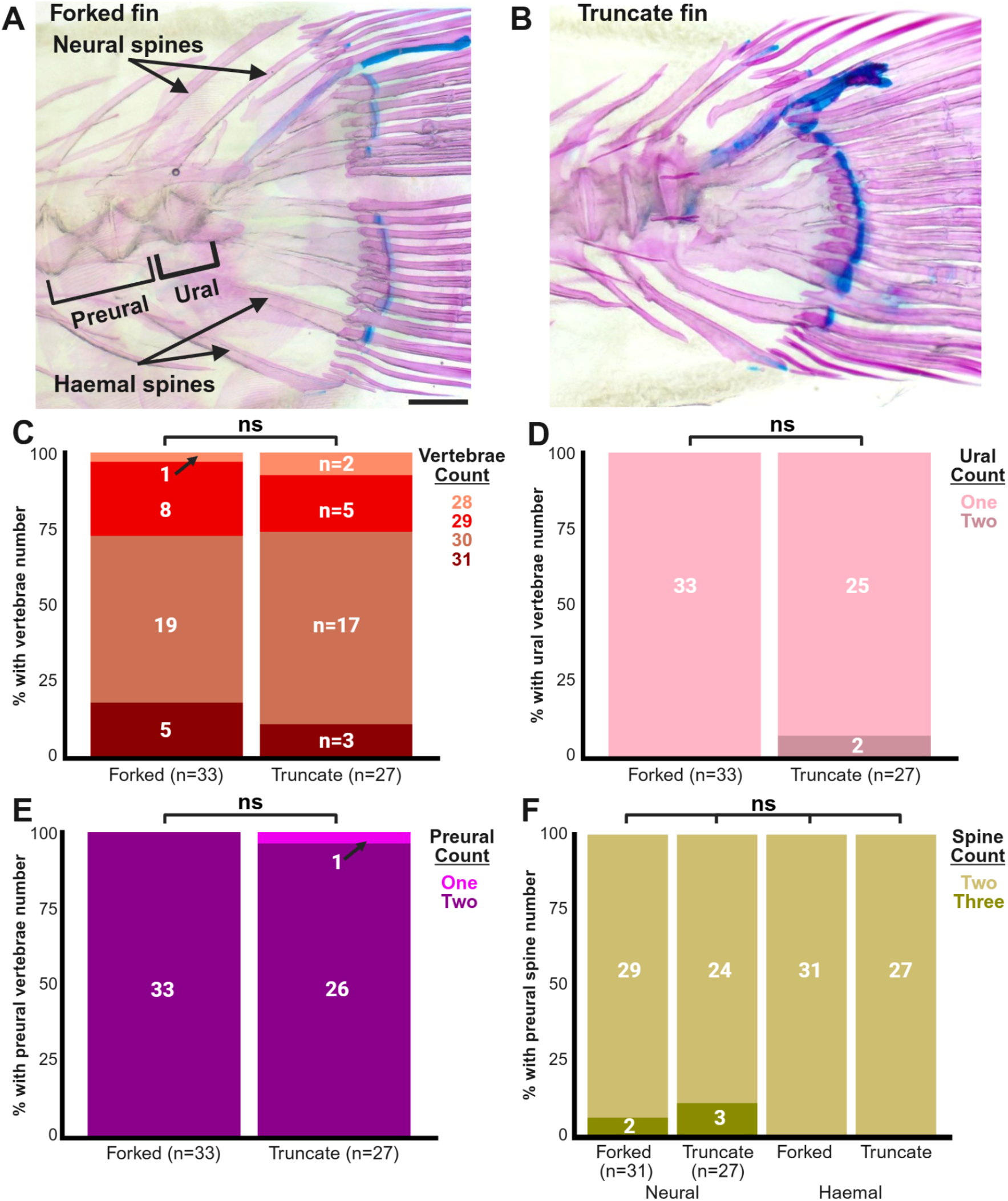
Vertebral and pleural phenotypes unaffected in truncate fin shape backgrounds. (*A-B*) Hypural complex of adult fins. Scale bar: 500 µm. There is no change in the number of total (*C*), ural (*D*) nor preural (*E*) vertebrae. Significance in the differences in means determined by Wilcoxon signed-rank test. (*F*) There is no change in the number of neural or haemal spines between WT forked or truncate caudal fins. Significance determined by two-way ANOVA followed by Tukey’s post hoc test.

**Fig. S7:**
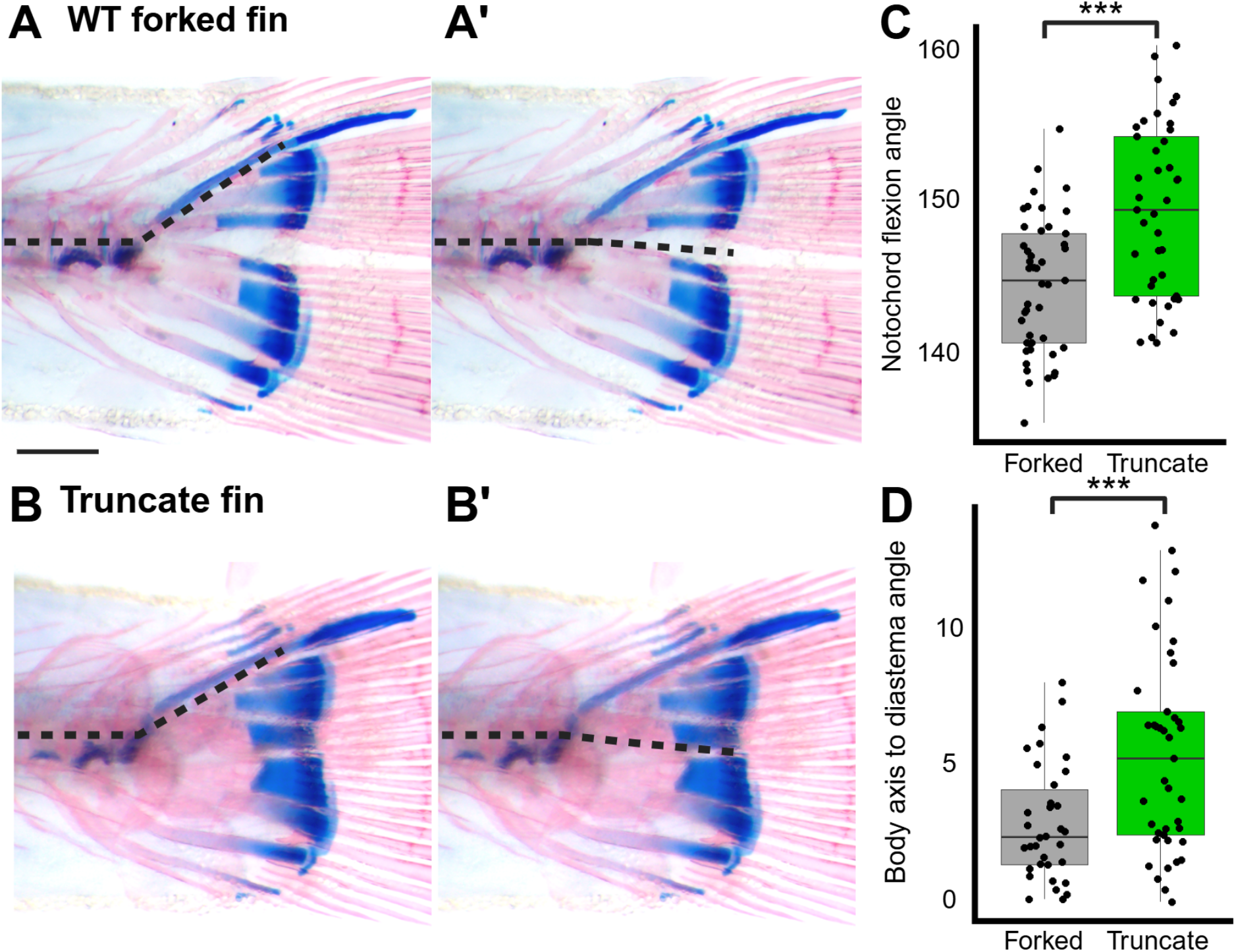
Notochord flexion and diastema-body axis alignment correlate with caudal fin shape. (A-B) Hypural complex of adult fins. Lines show angle of notochord flexion (A, B) and angle of body axis to diastema (A’, B’). Scale bar: 200 µM. Quantifications in (*C-D*), significance determined by Welch two-sample T-tests.

**Fig. S8:**
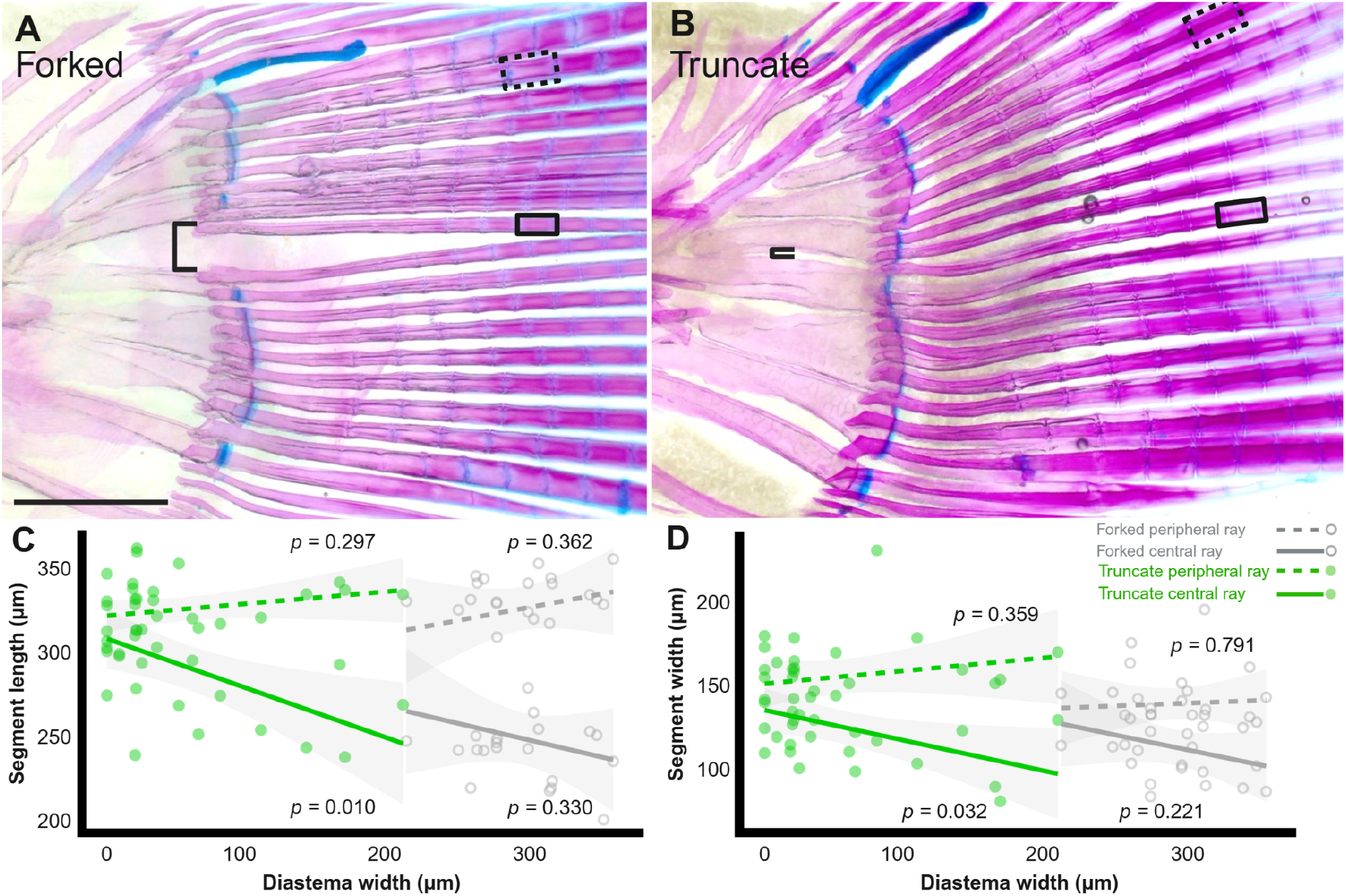
Central ray morphology correlates with hypural diastema aperture in truncate fins. In adult zebrafish caudal fins (*A-B;* cleared & stained), segment lengths (*C*) and widths (*D*) of the central fin rays in truncate fins (solid green line), but not peripheral (dotted green line), trend with the width of the hypural diastema (black bracket). Segment morphology in WT, forked fins do not correlate with hypural diastema width. Scale bar: 1 mm. Significance of relationship between segment morphology, fin shape, fin region and hypural diastema size is determined by linear-mixed effects model.

**Figure S9:**
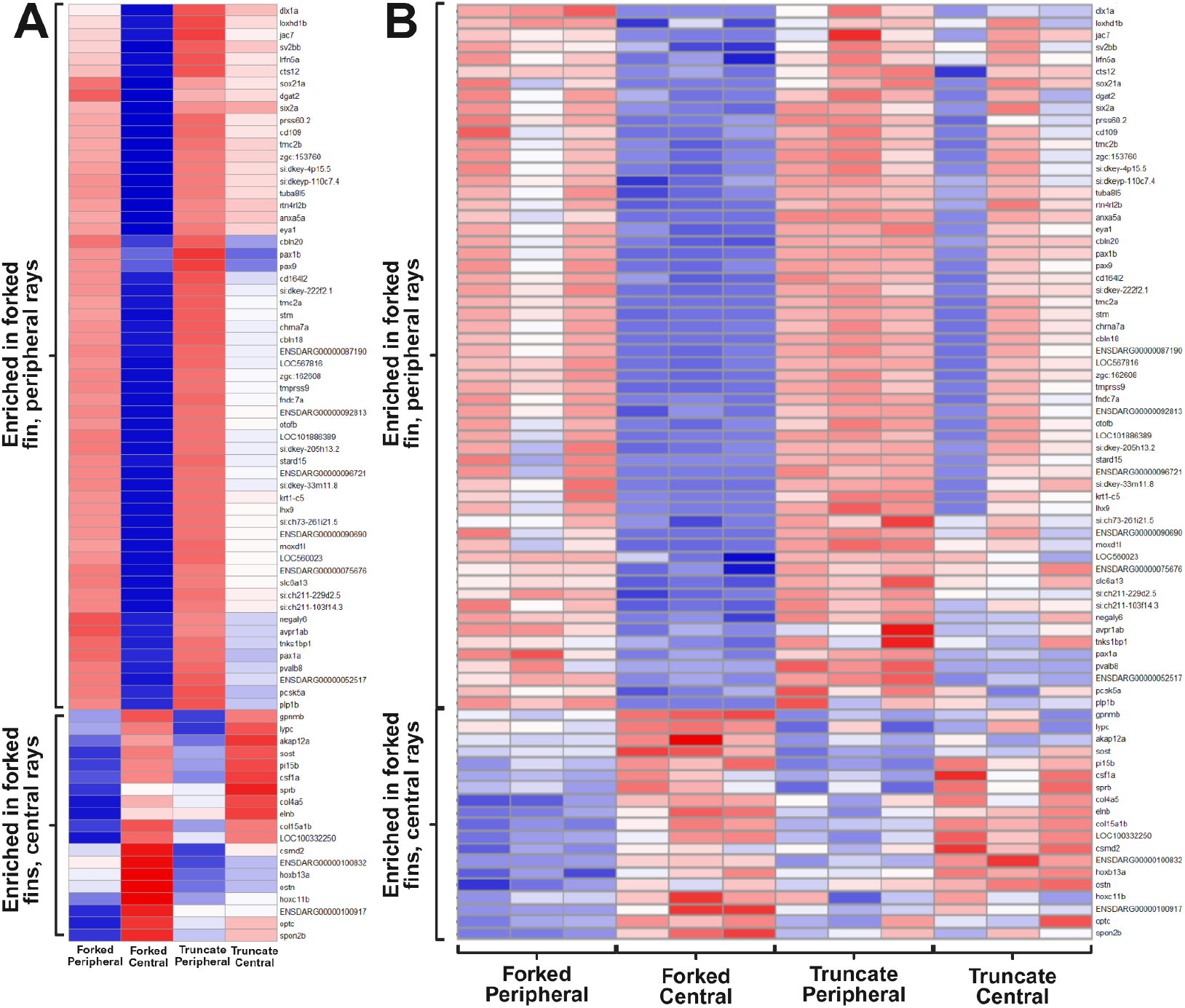
RNA sequencing comparisons. (*A*) Heatmap of differentially expressed differentially between central and peripheral regions of intact WT forked fins. *(B)* Expanded heatmaps showing the unpooled expression data for the genes shown in *(A)*, displaying all three individual replicates per condition.

**Fig. S10:**
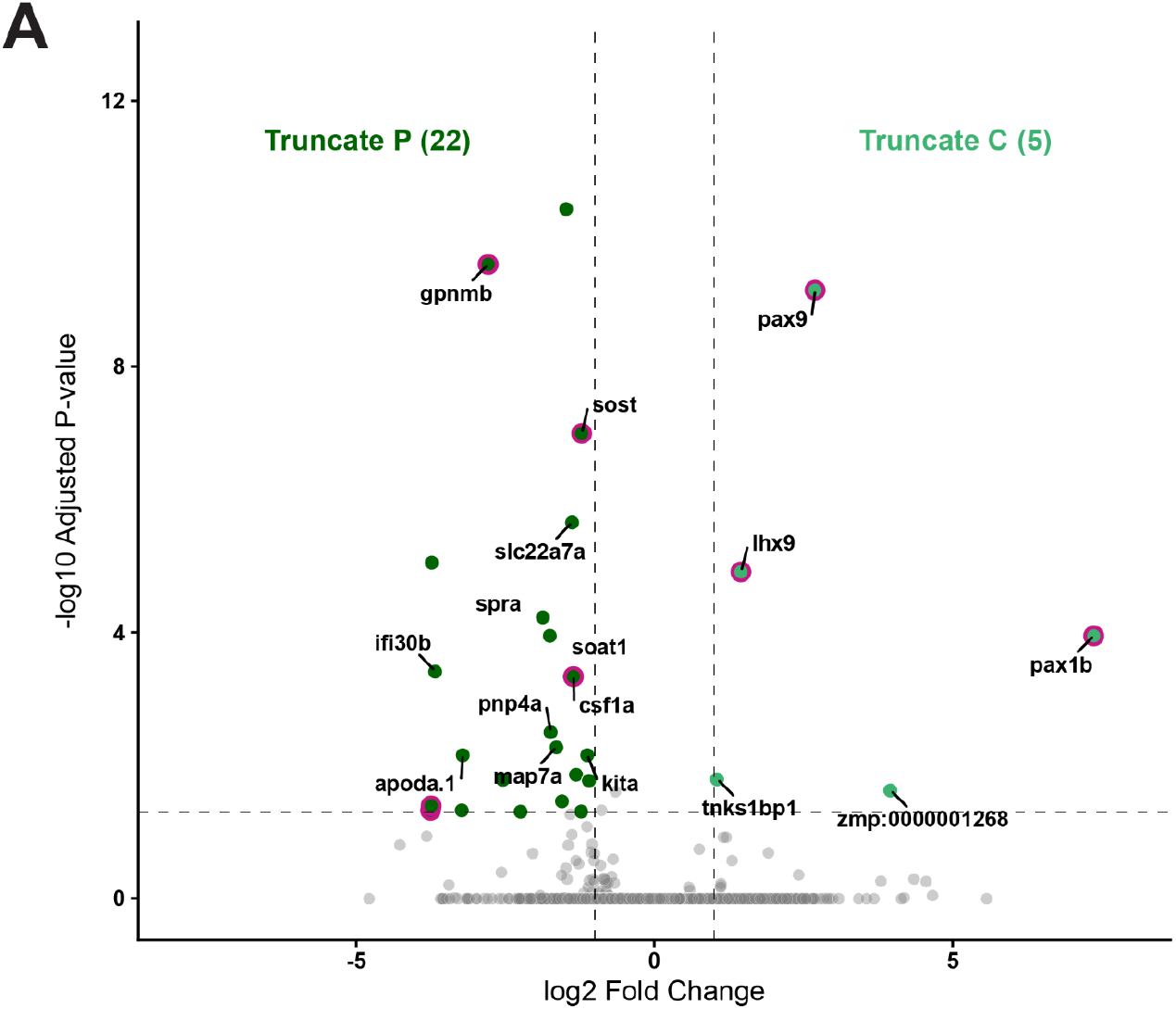
Truncate fins show relatively little expression difference between central and peripheral regions. (*A*) Volcano plot showing differentially expressed genes between central and peripheral fin rays in truncate fins.

**Fig. S11:**
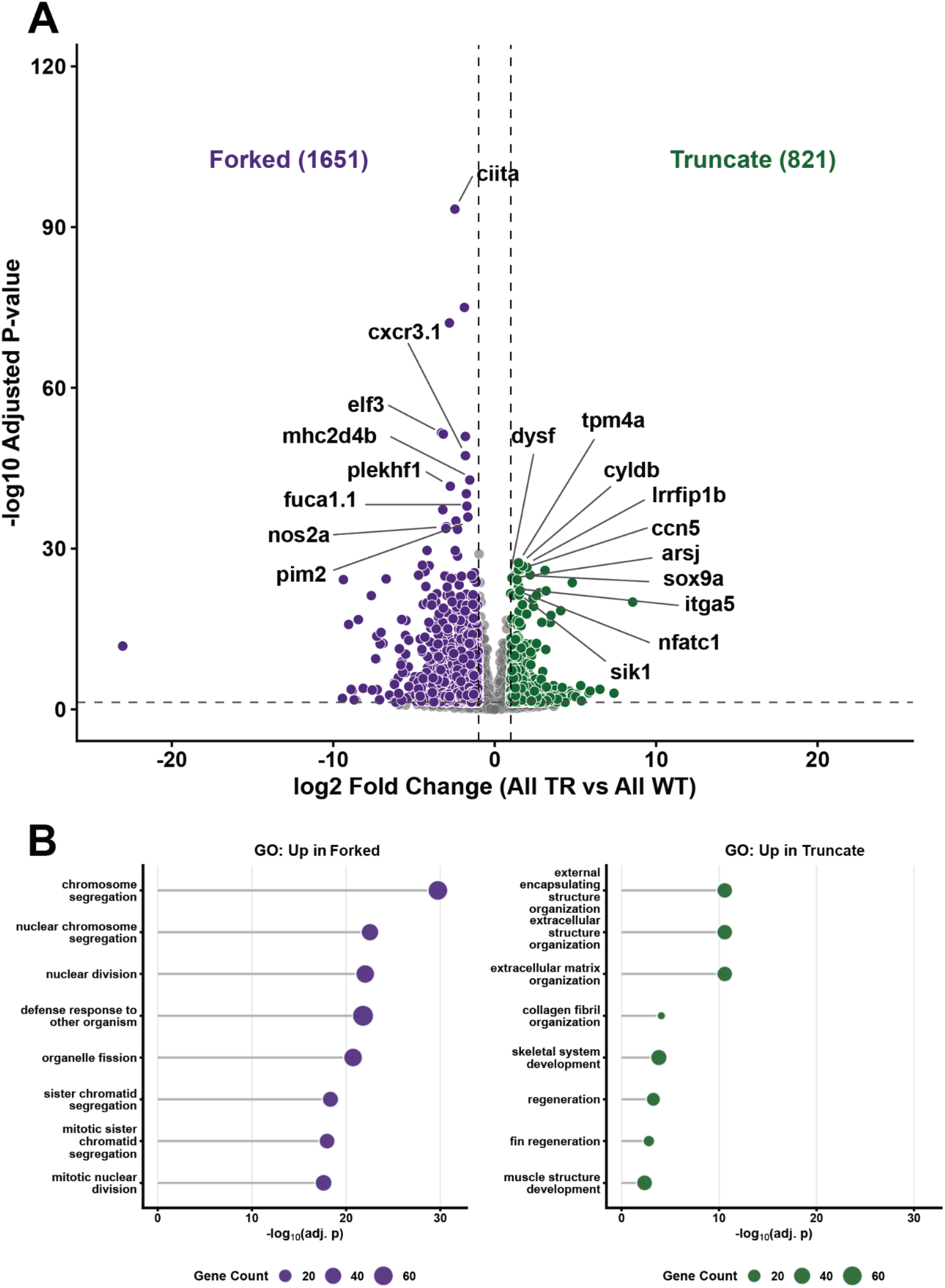
WT forked and truncate fins show unique patterns of enrichment. (*A*) Volcano plot showing differentially expressed genes between wild type forked and truncate (TR) fin rays, with central and peripheral samples pooled in each condition. *(B)* Gene ontology (GO) analysis showing enriched terms for the pooled WT forked and TR transcriptomes.

